# A hemoperfusion column selectively adsorbs LAP^+^ lymphocytes to improve anti-tumor immunity and survival of tumor-bearing rats

**DOI:** 10.1101/2024.05.28.596212

**Authors:** Kazuo Teramoto, Yuji Ueda, Ryosuke Murai, Kazumasa Ogasawara, Misako Nakayama, Hirohito Ishigaki, Yasushi Itoh

## Abstract

A decrease of immune suppressive cells in blood is thought to be one of the means to activate anti-tumor immunity that works as a treatment for cancers. We have developed an adsorbent that selectively adsorbs lymphocytes expressing latency-associated peptide (LAP), which include regulatory T cells (Tregs). The adsorbent, diethylenetriamine-conjugated polysulfone coated on polyethylene terephthalate fibers, was packed in a column for direct hemoperfusion (DHP). The therapeutic efficacy of DHP with the column was examined in rats carrying KDH-V liver cancer cells, in which LAP^+^ cells were increased in blood. After DHP, LAP^+^ T cells were decreased in peripheral blood, and a cytotoxic T-lymphocyte response against KDH-V cells was increased in tumor-bearing rats that had been immunized with X ray-irradiated KDH-V cells. Furthermore, the survival time of the rats was longer than that of rats without DHP. Thus, the removal of LAP^+^ T cells can potentially be applied to the treatment of cancer regardless of the origin since an increase in the number of LAP^+^ cells has been observed in the peripheral blood of various cancer patients.

## INTRODUCTION

The development of cancer treatments using new strategies is needed because progressive cancers have not been completely cured regardless of recent advances in cancer treatments. Immunotherapy using immune checkpoint inhibitors (ICIs) has recently been used as a treatment strategy in addition to surgical resection, chemotherapy, and radiotherapy. Inhibitors of PD-1/PD-L1 binding, which enhance T cell responses, have shown clinical efficacy for various tumors [1, 2]. Furthermore, anti-CTLA-4 antibody prolonged survival in the treatment of advanced melanoma [3]. However, not all cancer patients respond to treatment with ICIs and serious adverse events such as the development of autoimmune disease, which are called immune-related adverse effects (irAEs), are observed in some cases. Moreover, cancer recurrence despite continuous administration has been reported [4–6]. Thus, immune activation based on new methods is required for greater tumor reduction and improvement in the survival of patients.

Immune regulatory factors in tumor tissues and blood of hosts carrying tumors include immunosuppressive cytokines and cells. One of the immunosuppressive proteins is transforming growth factor (TGF)-β, which exists as a complex with latency-associated peptide (LAP) in plasma. The expression of TGF-β in a tumor was shown to be positively correlated with poor prognosis in patients with advanced gastric cancer [7]. Immunosuppressive cells in hosts carrying tumors were identified in CD25^+^CD4^+^ T cells, some of which are regulatory T cells (Treg) expressing the transcription factor Foxp3, called conventional Treg cells (cTreg) [8, 9]. On the other hand, it was reported that CD4^+^ T cells expressing a complex of TGF-β and LAP on the cell surface (LAP^+^CD4^+^ T cells) have a regulatory function [10]. LAP^+^ CD4^+^Foxp3^+^ T cells are a subgroup of Tregs and are enriched in blood and tumor tissue of patients with colorectal cancer [11]. LAP^+^CD4^+^ Foxp3^−^ T cells in colon cancer tumors showed a 50-fold higher immunosuppressive function than did cTreg cells [12]. Furthermore, LAP^+^CD8^+^T cells were also reported as immunosuppressive cells, suggesting that LAP^+^ cells are broadly immunosuppressive [13].

Increases of LAP^+^ cells were reported in tumor tissues and peripheral blood of various cancer patients including patients with colon, liver, and pancreas cancers [14–16]. Not only T cells and B cells but also immature myeloid cells include LAP^+^ cells that exhibit immunosuppressive properties [17–19]. Therefore, we thought that removing immunosuppressive proteins and regulatory cells would activate anti-tumor immunity. This speculation is supported by the results of our previous study showing that treatment of tumor-bearing rats with a column that adsorbs TGF-β resulted in prolonged survival [20] and the results of another study showing that reduction of LAP^+^ cells by administration of an anti-LAP antibody activated anti-tumor immunity in tumor-bearing mice [21]. Based on the results of those studies, we thought that the removal of LAP^+^ cells, a representative type of TGF-β-producing cells, would be more effective than removal of TGF-β proteins in tumor therapy.

In the present study, we examined the increase in the number of LAP^+^ cells in rats inoculated with liver cancer cells, KDH-V cells, and the survival of the cancer-bearing rats after direct hemoperfusion (DHP) with a LAP^+^ cell adsorbent column. The adsorbent column reduced the percentage of LAP^+^ cells in peripheral blood and upregulated cytotoxic T-lymphocyte (CTL) responses against tumor cells, resulting in improvement in survival of the rats.

## MATERIALS AND METHODS

### Preparation of tumor-bearing rats

This study was carried out in strict accordance with the Guidelines for the Husbandry and Management of Laboratory Animals of the Research Center for Animal Life Science at Shiga University of Medical Science and in strict accordance with Fundamental Guidelines for Proper Conduct of Animal Experiments and Related Activities in Academic Research Institutions under the jurisdiction of the Ministry of Education, Culture, Sports, Science and Technology, Japan. The protocol was approved by the Shiga University of Medical Science Animal Experiment Committee (Permit numbers: 2015-5-12, 2017-3-14, 2020-4-12). All procedures were performed by institutionally licensed researchers under anesthesia with medetomidine (0.15 mg/kg), midazolam (2.0 mg/kg), and butorphanol tartrate (2.5 mg/kg) or sodium pentobarbital (200 mg/kg). The rats were monitored daily during the study. The rats were euthanized at a humane endpoint when the tumor diameter reached 3 cm. The other rats were observed as long as 80 days after tumor inoculation and thereafter were euthanized for sample collection.

We previously established a tumor-bearing rat model by inoculating syngeneic WKAH/Hkm rats with transplantable KDH-8 cells derived from chemically induced liver cancer developed at Hokkaido University [22], and the mean survival time of the rats was 55 days. Most of the KDH-8 cells are adherent to the polystyrene dishes when cultured in RPMI1640 medium (Nacalai Tesque, Kyoto, Japan) supplemented with 10% fetal calf serum (FCS). After the floating cells were cultured repeatedly, completely floating cells were obtained and the cells were named KDH-V cells. KDH-V cells (3 × 10^5^ cells) were dispersed in phosphate-buffered saline (PBS) and injected subcutaneously into the backs of WKAH/Hkm rats to prepare tumor-bearing rats. WKAH/Hkm rats were obtained from Japan SLC Inc. (Hamamatsu, Japan) and were reared at the Research Center of Animal Life Science, Shiga University of Medical Science.

### Preparation of rats immunized with irradiated KDH-V cells

To induce KDH-V-specific immune responses in WKAH/Hkm rats, KDH-V cells were irradiated with 10,000 R of X-ray using an irradiator (MBR1520R; Hitachi Medico, Ltd., Tokyo, Japan) at the Central Research Laboratory, Shiga University of Medical Science. Irradiated KDH-V cells (X-KDH, 1 × 10^7^ cells) in 0.5 mL of PBS were inoculated subcutaneously in the backs of 4- to 8-week-old WKAH/Hkm rats. After 3 weeks or more, the immunized rats were subjected to the following experiments.

### Analysis of blood cells

Blood (0.5 mL) was collected from the subclavian vein using a 23G × 1/4″ injection needle under anesthesia with medetomidine, midazolam, and butorphanol tartrate. The blood cell composition was measured using VetScan HMII (Abaxis Inc., Union City, CA).

For flow cytometric analysis, an antibody cocktail shown in S1 Table was added to 90 μL of whole blood cells and the cells were incubated at room temperature for 30 min, and then the cells in a washing buffer were centrifuged at 1,400 rpm for 5 min. Thereafter, red blood cells were lysed in FACS™ Lysing Solution (BD Biosciences, Franklin Lakes, NJ). More than 50,000 cells were analyzed with the CytoFlex S flow cytometer (Beckman Coulter, Inc., Brea, CA).

### *In vitro* CTL assay

Effector cells were prepared from homogenized spleens after purification using the Lymphocyte Separation Medium (density: 1.077) (Wako Pure Chemicals, Osaka, Japan). For target cells, KDH-V cells (1 × 10^7^ cells/mL in PBS) were labeled with carboxyfluorescein diacetate succinimidyl ester (CFDA-SE) (Dojindo Laboratories, Kumamoto, Japan) for 10 min at 37°C. For a CTL assay, effector cells and target cells were mixed and cultured at E/T ratios of 100, 50, 25 and 12.5 in 48-well plates for 22 h at 37°C (n = 4). After culture, propidium iodide (PI) was added and 1 × 10^5^ cells were acquired for the flow cytometer assay. PI-positive cells were recognized as dead KDH-V cells among the carboxyfluorescein succinimidyl ester (CFSE)-positive cells.

### Detection of interferon-gamma (INF-γ) production using ELISpot

For an ELISpot assay, the IFN-γ EL-585 kit (R&D systems, Minneapolis, MN) was used according to the manufacturer’s instruction. Purified peripheral blood mononuclear cells (PBMC) were cultured for 20 h in ELISpot plates and then colored. IFN-γ-positive spots were counted using the ImmunoSpot S6 ULTRA Basic Analyzer (Cellular Technology Limited, Cleveland, OH) (n = 6 to 8 wells).

### Preparation of hemoperfusion columns

According to the chemical formula shown in Fig 1A, chloro-acetamidomethylated polysulfone was prepared by amidomethylation of polysulfone (Mn ∼22,000; Sigma-Aldrich) with N-hydroxymethyl-2-chloroacetamide. The obtained polymer was coated on non-woven fabric fibers made of polyethylene terephthalate (PET) and reacted with diethylenetriamine to produce an adsorbent. A cylindrical column with an inner diameter of 1 cm and a length of 2 cm was filled with 0.25 g of the resulting adsorbent for DHP.

**Fig 1.**
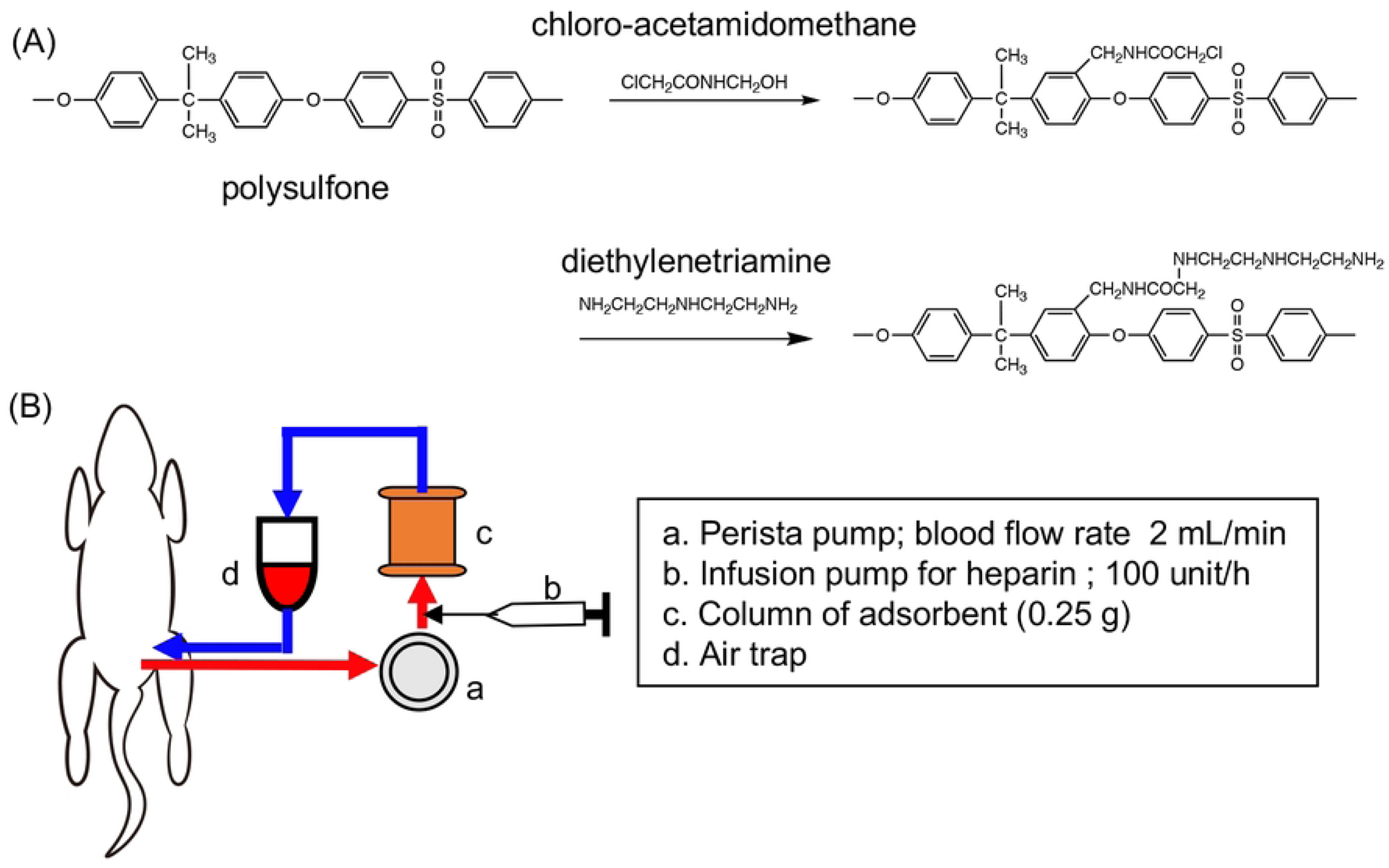
LAP^+^ cell adsorbent material and direct hemoperfusion. (A) Chemical reaction for preparation of the adsorbent. (B) Scheme of direct hemoperfusion. Blood was drawn from the femoral artery and passed through the column driven by a Perista pump to return to a femoral vein.

### Procedure for DHP

WKAH/Hkm rats were anesthetized by subcutaneous injection of sodium pentobarbital (200 mg/kg) (Fig 1B). The groin skin on one side was incised to expose the femoral artery and vein, and then two 24G×3/4″ Surflo F&F needles (Terumo, Tokyo, Japan) were inserted into the artery and vein to connect to an extension tube X1-50 (Top Ltd., Tokyo Japan). Saline containing 200 units of heparin (2 mL, AY Pharma, Tokyo, Japan) was administered through the vein. Femoral arterial blood was led to a three-way stopcock, a 2 × 4 mmφ silicon tube that was set in a microtube pump MP2000 (Tokyo Rikakikai Co., Ltd., Tokyo, Japan), the column, an air trap, and a three-way stopcock to return the blood to the femoral vein. Immediately before extracorporeal circulation, the circuit was washed with 40 mL of saline containing 200 units of heparin. DHP was performed for 1 h. During the circulation, heparin was continuously infused at a rate of 100 units/h using a mini-syringe pump (Terumo Corp., Tokyo, Japan). After DHP, the catheter was removed from the artery and 4 mL of saline was flushed to return the blood. The skin was sutured after returning the blood and removing the catheter from the vein.

## Results

### Changes in white blood cells in a KDH-V rat tumor model

We established a rat tumor model to evaluate the efficacy of treatment with a hemoperfusion column. After 3 × 10^5^ KDH-V cells were subcutaneously inoculated in the backs of naïve rats (unimmunized rats) and rats immunized previously with the irradiated KDH-V cells (immunized rats), the tumor became palpable on day 7 and thereafter the tumor gradually increased in size over time (Fig 2A). Tumor sizes reached the humane endpoint 21 days and 35 days after tumor inoculation in unimmunized and immunized rats, respectively. Thus, immunization with irradiated tumor cells prolonged the survival time of the rats inoculated with KDH-V cells.

**Fig 2.**
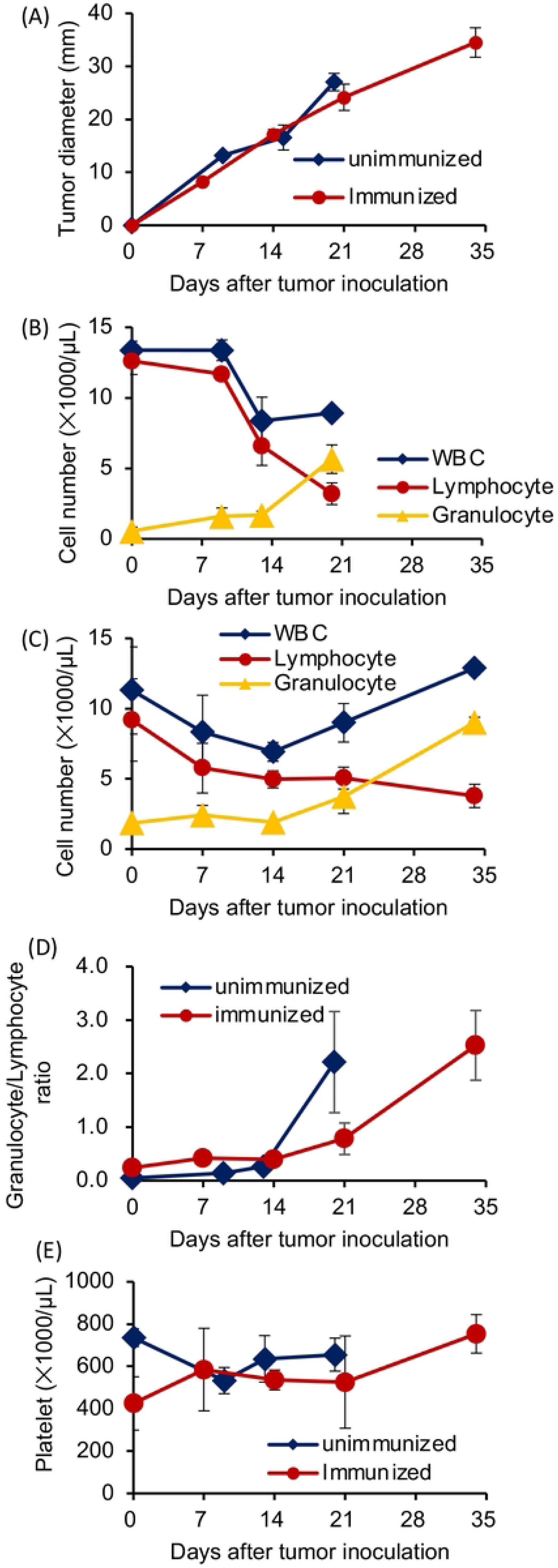
Peripheral blood cell changes in rats carrying KDH-V cells. Unimmunized rats (n = 3) and rats immunized with irradiated KDH-V cells (n = 3) were subcutaneously injected with 3 × 10^5^ KDH-V cells into the backs. Blood (0.5 mL) was collected from the subclavian vein every week. (A) Tumor growth size is shown as an average of the larger and smaller diameters. (B, C) White blood cell counts in unimmunized rats (B) and in immunized rats (C). (D) The granulocyte/lymphocyte ratio in peripheral blood of rats was calculated on the basis of results of (B) and (C). (E) Platelet counts in unimmunized and immunized rats.

Peripheral blood cells in rats bearing KDH-V cells were examined. In unimmunized rats, the mean white blood cell count did not change until day 9 and was decreased on day 13. The mean lymphocyte count was decreased on day 13, whereas the mean granulocyte count was increased on day 21 (Fig 2B). On the other hand, in immunized rats, the mean white blood cell counts were gradually decreased until day 14 and were increased on days 20 and 34. Moreover, the mean lymphocyte counts in immunized rats were gradually decreased, whereas the mean granulocyte counts were increased after day 21 (Fig 2C). Granulocyte/lymphocyte ratios were increased on day 20 and day 34 in unimmunized rats and immunized rats, respectively (Fig 2D). The mean platelet counts in unimmunized rats were slightly decreased on day 10 and increased on day 13, whereas the mean platelet count in immunized rats was increased on day 34 (Fig 2E). Thus, the number of peripheral white blood cells in the tumor-bearing rats changed with tumor growth.

### LAP-positive cells in peripheral blood of rats carrying KDH-V cells

We determined the percentage of cells expressing LAP (LAP^+^ cells) on the cell surface by flow cytometry. Lymphocytes (P2), monocytes (P3), and granulocytes (P4) in CD45^+^ cells (P1) were defined in FSC/SSC dot plots (Fig 3A, B). Among CD3^+^ cells (P5) (Fig 3C), CD4^+^ T and CD8^+^ T cells were identified by their respective surface antigens (Fig 3D). CD45RA^+^ and CD3^−^ cells were considered as B cells (Fig 3E). Thereafter, the percentages of LAP^+^ cells (red dots) were determined in comparison with control staining (blue dots) (Fig 3 E-I). The percentages of LAP^+^ cells in granulocyte marker-positive cells in the P4 and P3 fractions were defined as granulocytes and monocytes, respectively (Fig 3H, I).

**Fig 3.**
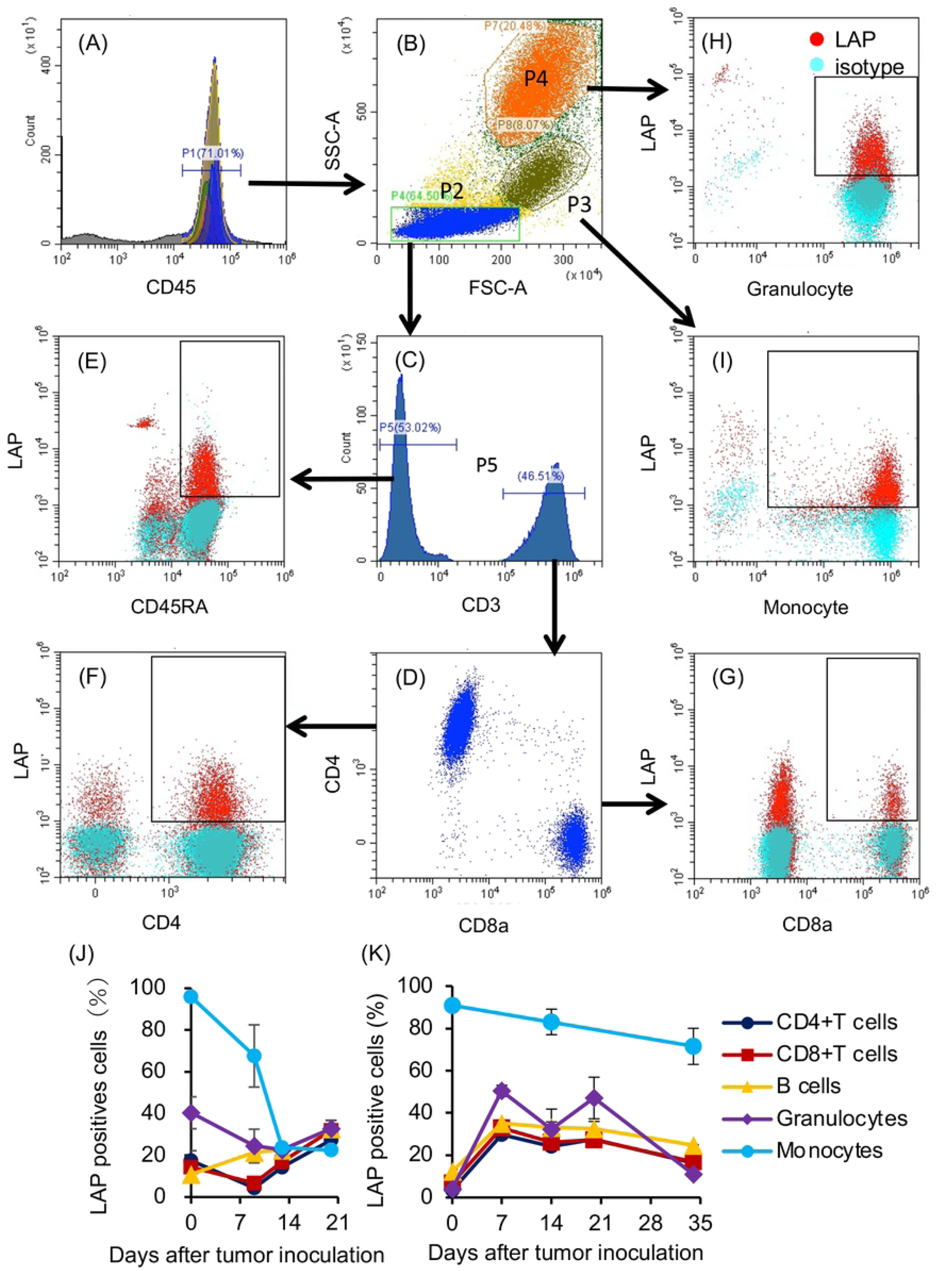
Percentages of LAP^+^ cells in peripheral blood cells of rats carrying KDH-V cells. A representative flow cytometry profile is shown as a gating example. (A) White blood cells were gated as CD45-positive cells and then (B) lymphocytes (P2), monocytes (P3), and granulocytes (P4) were separated in an FSC/SSC plot. (C, D) CD4^+^ T cells and CD8^+^ T cells were identified among CD3^+^ cells and (E) B cells were identified as CD45RA^+^ cells among CD3^−^ cells. The percentages of LAP^+^ cells in CD4^+^ T cells, CD8^+^ T cells, B cells, granulocytes, and monocytes were determined (E–I). The means and standard deviations of percentages of LAP^+^ cells in each population were calculated in (J) unimmunized rats (n = 3) and (K) immunized rats (n = 3).

We examined changes in LAP ^+^ cells in the tumor-bearing rats. The mean percentages of LAP^+^ cells in CD4^+^ T cells, CD8^+^ T cells, and granulocytes were decreased on day 9 (4.3%, 6.7%, and 24.4%, respectively) compared to the percentages on day 0 before tumor inoculation (17.4%, 14.3%, and 40.4%, respectively) and they were increased on day 20 in unimmunized rats (27.3%, 31.9%, and 32.7%, respectively) (Fig 3J). The mean percentages of LAP^+^ B cells were increased on day 9 (21.5%) and day 20 (32.7%) compared to the mean percentage on day 0 (10.9%), whereas the mean percentages of monocytes, which were mostly positive for LAP before tumor inoculation, were decreased on day 9 (67.6%) and day 20 (22.4%) compared to the mean percentage on day 0 (95.9%).

The percentages of LAP^+^ cells in immunized rats showed a different tendency from that in unimmunized rats. The mean percentages of LAP^+^ cells in CD4^+^ T cells, CD8^+^ T cells, B cells, and granulocytes were increased on day 7 after tumor inoculation (29.7%, 32.9%, 35.1%, and 50.4 %, respectively) and they were decreased on day 34 (16.4%, 16.9%, 24.7%, and 10.9%, respectively) (Fig 3K). On the other hand, the percentage of LAP^+^ cells in monocytes gradually decreased to 71.5% on day 34. Thus, the percentages of LAP^+^ cells in lymphocytes and granulocytes showed increases in early time points but decreased in the final stage after tumor inoculation in immunized rats.

### Improvement of anti-tumor immunity in tumor-bearing rats by DHP

Since removal of TGF-β using a direct hemoperfusion (DHP) fiber column prolonged survival in a KDH-8 rat cancer model in our previous study (20), we tried to remove LAP^+^ cells, which are representative TGF-β-producing cells, in the present study. After coating the surface of the PET fiber with chloroacetamidomethylated polysulfone, a diethylenetriamine group was introduced to provide functional groups to prepare an adsorbent, which was packed in a column (Fig 1). Twelve days after inoculation of KDH-V cells into unimmunized rats, tumor-bearing rats were subjected to DHP (Fig 4A). Blood cell counts before and after circulation showed a slight increase in the lymphocyte count (Fig 4B), whereas granulocytes and platelets decreased to 45% and 36% of the counts before circulation, respectively (Fig 4B and C). The levels of reduction of LAP^+^ cells by DHP were 40% for CD4^+^ T cells and 41% for CD8^+^ T cells (Fig 4D).

**Fig 4.**
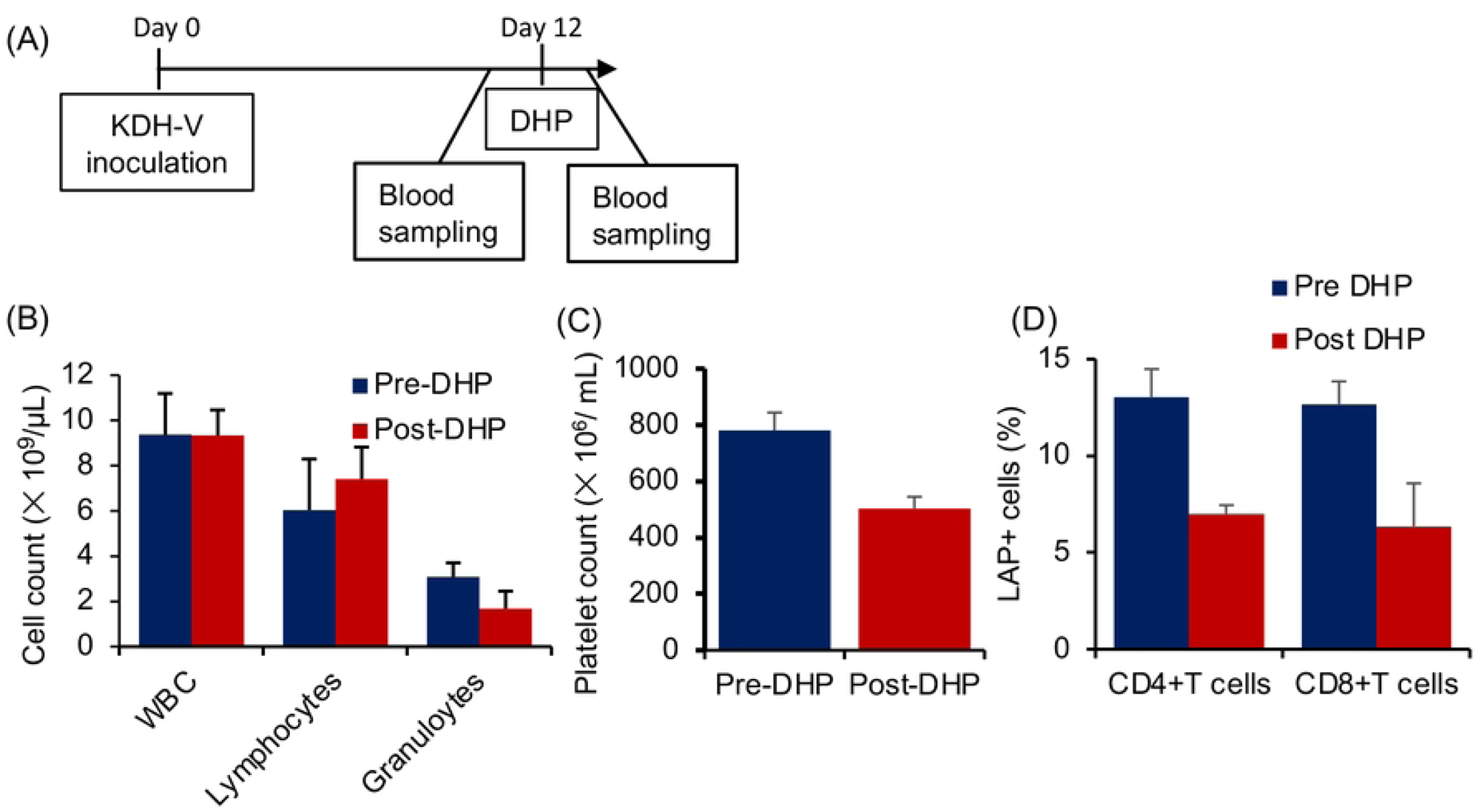
Effects of the DHP column on LAP^+^ cells in peripheral blood of unimmunized rats carrying KDH-V cells. (A) DHP was performed 12 days after subcutaneous inoculation of KDH-V cells and blood was collected before and after DHP. (B) Means and standard deviations of total white blood cell, lymphocyte, and granulocyte counts before and after DHP (n = 3). (C) Means and standard deviations of platelet counts before and after DHP (n = 3). (D) Means and standard deviations of the percentages of LAP^+^ cells in CD4^+^ T cells and CD8^+^ T cells (n = 8).

To confirm whether the anti-tumor activity was increased by the DHP, 4 unimmunized rats were inoculated with KDH-V cells, and then 2 rats underwent DHP to examine blood cells, tumor-infiltrating lymphocytes (TIL), and CTLs in the spleen (Fig 5A). In the blood 8 days after DHP treatment (day 15), the mean percentages of LAP^+^ cells in CD4^+^ T cells and CD8^+^T cells of DHP-treated rats were 33% and 32%, respectively, lower than those in untreated rats (Fig 5B). In addition, the percentages of LAP^+^ cells in CD4^+^ T cells, CD8^+^ T cells, and granulocytes in tumor tissues (TIL) were 52%, 62%, and 33% lower in DHP-treated rats than in untreated rats, respectively (Fig 5C). Moreover, CTL activity of the splenocytes against KDH-V cells was higher in DHP-treated rats than in untreated rats (Fig 5D). Furthermore, after the immunized rats underwent hemoperfusion with a column on day 11 after tumor inoculation, the CTL activity was examined 3 days after DHP treatment (day 14) (Fig 5E). The CTL activity against KDH-V cells was higher in splenocytes from DHP-treated immunized rats than in those from untreated immunized rats (Fig 5F). Thus, CTL activity specific for KDH-V cells was upregulated after DHP treatment in both unimmunized and immunized rats.

**Figure 5.**
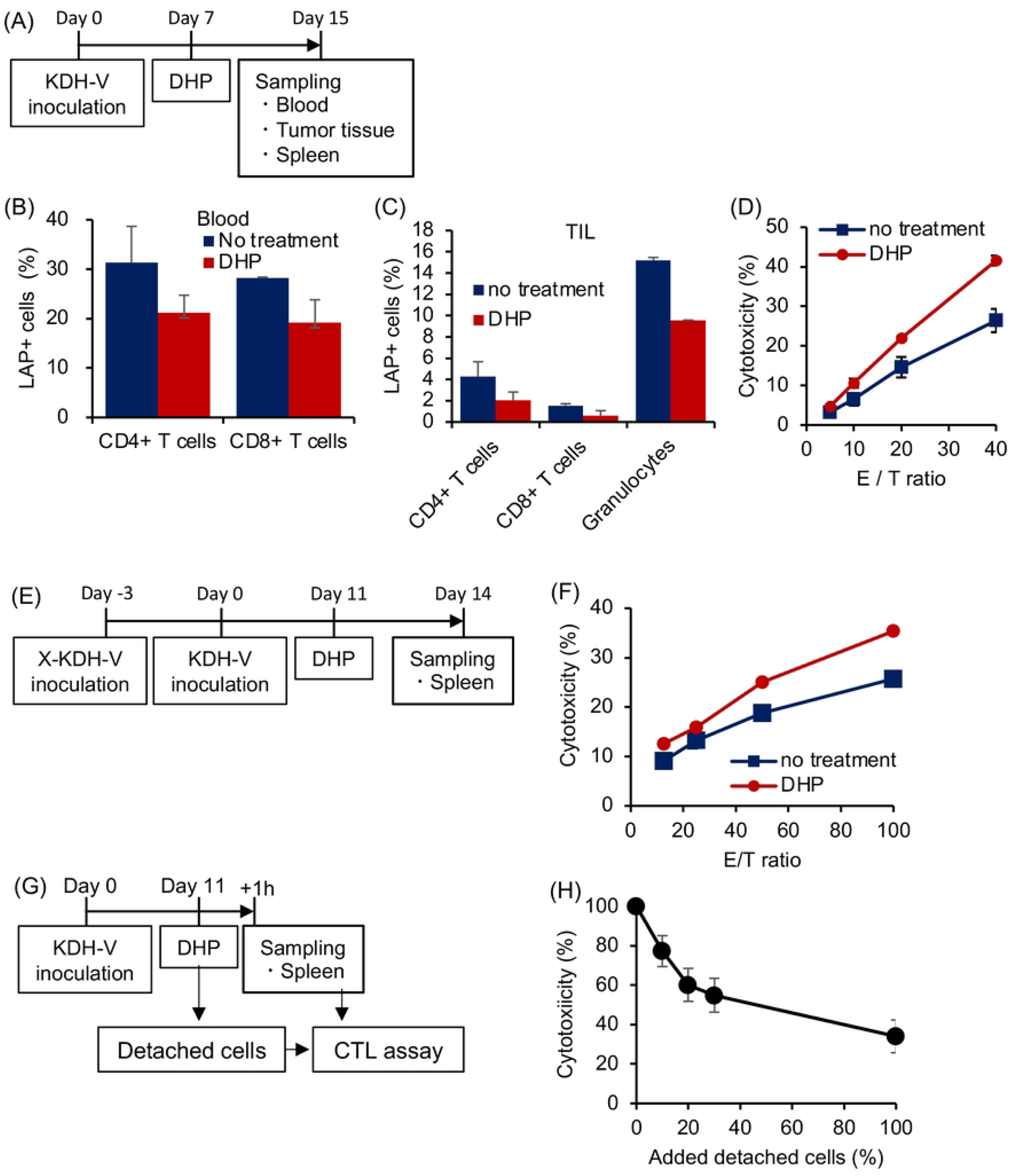
LAP^+^ T cells in tumor tissues and cytotoxic T-lymphocyte activity before and after DHP. (A) DHP was performed 7 days after subcutaneous inoculation of KDH-V cells, and blood, tumor tissues, and spleens were collected 8 days after DHP. (B, C) Means and standard deviations of percentages of LAP^+^ cells in CD4^+^ T cells and CD8^+^ T cells in peripheral blood (B) and tumor tissues (C) were calculated. (D) CTL activity of spleen cells against KDH-V cells was analyzed using a flow cytometer. The percentage of cytotoxicity was measured at different effector/target ratios (n = 3). (E, F) CTL activity against KDH-V cells of spleen cells from immunized rats was measured (n = 3). (G, H) Suppressive activity of cells adsorbed in the DHP column was determined by addition of detached cells in the CTL activity culture. Means and standard deviations of percent cytotoxicity were calculated in the different numbers of detached cells (n = 3).

In addition, the immunosuppressive function of cells adsorbed to the column was examined. The adsorbed cells were detached from the column with PBS after hemoperfusion on day 11 after tumor inoculation. When the collected cells were mixed with splenocytes from the same unimmunized rat (Fig 5G), the CTL activity decreased depending on the number of adsorbed cells added (Fig 5H) (n = 3). This result indicates that the DHP column removes immunosuppressive cells from rats carrying a tumor.

We further examined the duration of immune upregulation after the DHP by an ELISpot assay. After immunized rats had been inoculated with KDH-V cells into their backs, they were subjected to DHP on day 12 or day 14. Peripheral blood was collected from the rats on day 17 (S1 Fig A). IFN-γ-producing cells specific for tumor antigen were detected in an ELISpot assay. The number of IFN-γ-producing spots was larger in blood of rats that had undergone DHP than in blood of untreated rats on 5 and 3 days after the DHP (S1 Fig B). Similar results were obtained in rats on 1 and 3 days after DHP (S1 Fig C-H). Thus, the upregulation of INF-γ-producing cells continued for at least 5 days after DHP.

### Survival time of rats carrying KDH-V cells after DHP treatment

Finally, we examined the effects of the LAP^+^ cell adsorbent column on survival of rats carrying KDH-V cells. Immunized rats were inoculated with KDH-V cells and then DHP treatment was performed for 60 min on days 5–7 (Fig 6A). Rats in the DHP-treated group survived significantly longer than did rats in the non-treated group (log-rank test, p = 0.005) (Fig 6B). On the other hand, in unimmunized rats, the DHP treatment did not extend the survival period (Fig 6C, D). These results suggest that the LAP^+^ cell adsorbent column upregulates the tumor-specific CTL response to reduce tumor progression.

**Fig 6.**
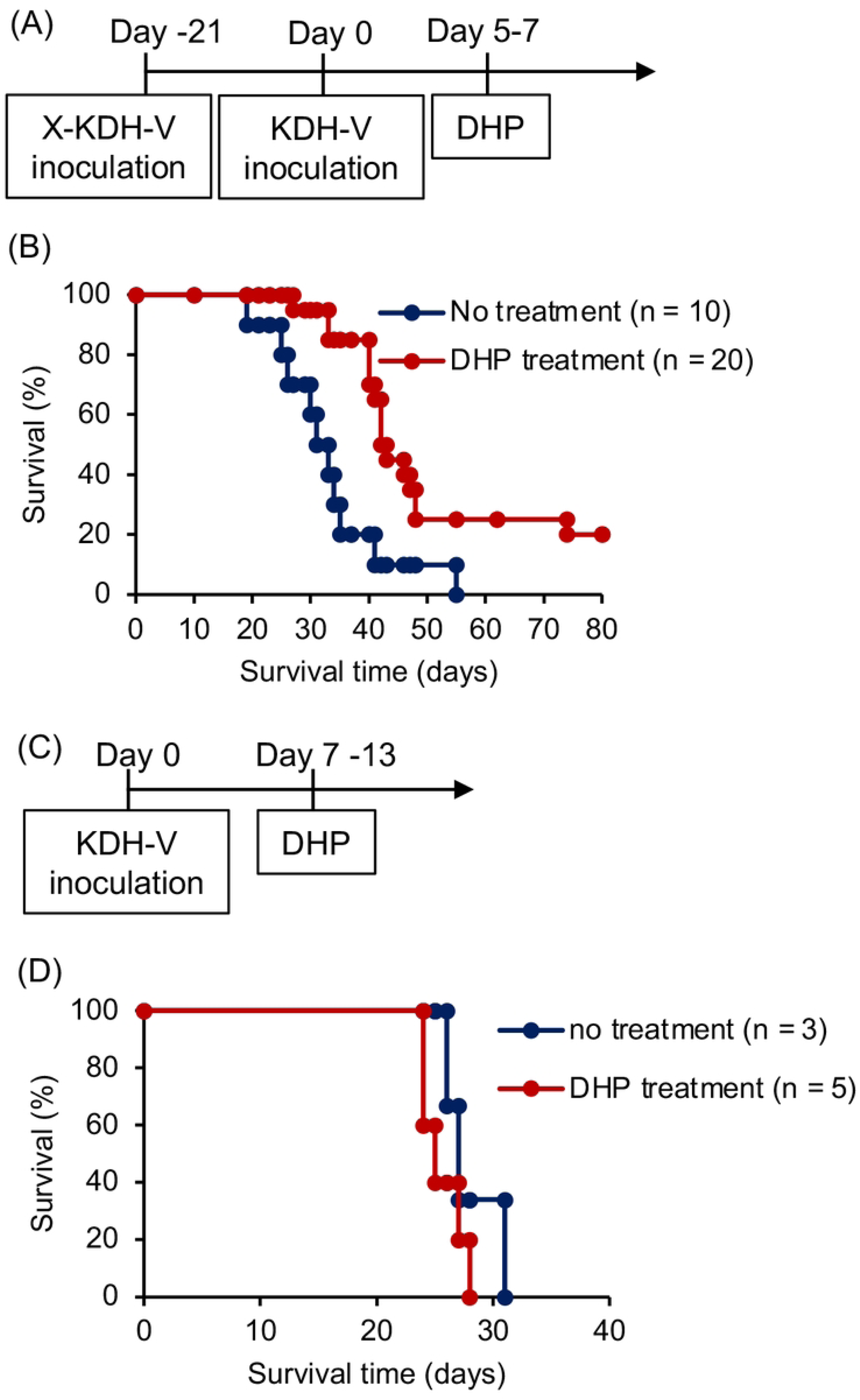
Survival of rats carrying KDH-V cells after DHP treatment that removed LAP^+^ cells. KDH-V cells were subcutaneously inoculated in the backs of immunized rats (A) with or (C) without immunization with irradiated KDH-V cells. Immunized rats were treated with DHP 5 to 7 days after tumor inoculation (B) and unimmunized rats were treated 7 to 13 days after tumor inoculation (D). (B, D) Survival rates of DHP-treated rats (red line) and non-treated rats (blue line).

## DISCUSSION

We have newly developed a column that adsorbs LAP^+^ cells in peripheral blood and that extends the survival period of tumor-bearing rats by DHP. The cancer model rat used in the present study showed a high percentage of LAP^+^ T cells in peripheral blood. After treatment with the LAP^+^ cell-adsorbing column, reduction of LAP^+^ T cells was observed in tumor tissue of the cancer rats as well as in peripheral blood. The CTL activity of splenocytes in the tumor-bearing rats treated with DHP was higher than that in the tumor-bearing rats without DHP. The cells adsorbed in the column were immunosuppressive because they inhibited CTL activity dose-dependently in the cell culture. Thus, reduction of LAP^+^ cells with DHP is thought to have the potential for cancer immunotherapy.

In the present study, reduction of LAP^+^ cells in peripheral blood activated an anti-tumor immune response and extended the survival period of rats carrying the liver cancer cells. In our previous study, we revealed that removal of the immunosuppressive protein TGF-β by using a TGF-β-adsorbing column improved the survival of tumor-bearing rats [20]. Therefore, we further developed the column to remove cells producing TGF-β, i.e., LAP^+^ cells, because we postulated that the removal of LAP^+^ cells would be more effective than removal of TGF-β for enhancing anti-tumor acquired immunity. The percentages of LAP^+^ cells in blood of the column-treated rats were lower than those in blood of the untreated rats 8 days after DHP. Moreover, the CTL activity of splenocytes in the column-treated rats was higher than that in the untreated rats 8 days after DHP. Therefore, the CTL activation effect was sustained for at least 8 days after DHP treatment.

The effects of DHP on amelioration of the survival period and rate were significant in immunized rats but not in unimmunized rats. This result indicates that LAP^+^ T cells affect acquired immune responses including a CTL response. Priming and expansion of CTLs specific for tumor antigens follows activation of innate immunity and antigen presentation. Therefore, immunization with irradiated KDH-V cells might allow the priming of CTL responses specific for tumor antigen and reduction of LAP^+^ cells might help the expansion and effector function of CTLs, effects of which were not seen in the unimmunized rats without priming of CTLs specific for tumor antigen. Because the KDH-8 tumor growth, which might allow the priming of CTLs, was slower than that of KDH-V, it was thought that DHP absorbing TGF-β showed anti-tumor effects in the previous study [20]. Therefore, the present model with rapid tumor growth and preimmunization ensures CTL priming in rats before the DHP treatment.

The removal of LAP^+^ cells using DHP has advantages in comparison with other immunotherapies. Administration of an anti-LAP antibody improved anti-tumor immunity of mice in a previous study [21] in which the anti-LAP antibody was injected every three days for one month after tumor implantation for elongation of survival time. In addition, the effect of antibody therapy was gradually attenuated after repeated administration as reported in ICIs [4–6]. On the other hand, hemoperfusion therapy using the DHP column was effective for improving the survival of tumor-bearing rats with only one DHP treatment, and enhancement of the CTL response by the column was effective for at least 8 days. These are advantages of hemoperfusion therapy in anti-tumor therapy. Repeated column treatment may be considered to further improve the efficacy of the hemoperfusion therapy. One of the concerns arising from hemoperfusion therapy is reduction of platelets, which might induce a bleeding tendency. In our additional study, the platelet counts after DHP were above 300,000/μL and they returned to the basal level 5 to 7 days after hemoperfusion therapy in the unimmunized rat model (S2 Fig). Therefore, bleeding due to a low platelet count is thought to be unlikely if the interval between hemoperfusion treatments is 7 days.

It is thought that removal of LAP^+^ cells can be applied for patients with various cancers. LAP^+^ T cells, which include some Foxp3^+^ T cells, increase in patients with colorectal cancer, non-muscle-invasive urinary bladder cancer, and other cancers [14–16, 23, 24]. In addition, the percentage of FoxP3^+^ Tregs in patients with non-muscle-invasive urinary bladder cancer was shown to be correlated with risk stratification and recurrent-free survival [23]. The percentage of LAP^+^CD4^+^ T cells in peripheral blood of hepatocellular carcinoma patients was reported to be correlated with tumor size [24]. These results suggest that LAP^+^CD4^+^ T cells and FoxP3^+^ Tregs migrate from peripheral blood to tumor tissues or circulate in the tissues. Therefore, the removal of LAP^+^ cells could be applied for treatment of patients with various cancers who have a high percentage of LAP^+^ cells in peripheral blood.

In the present study, we showed the efficacy of hemoperfusion column therapy in a rat cancer model. The removal of LAP^+^ cells could be applied to various tumors since an increase of LAP^+^ cells has been reported in other cancers [13–16]. Since the column removes regulatory lymphocytes, it consequently has an ameliorative effect on immune responses regardless of the type of cancer. Thus, removal of LAP^+^ cells from cancer patients may be an alternative immune therapy to ICIs.

## Supporting information

**S1 Table. Combination of antibodies to detect surface antigens**

**S1 Fig. Duration of the effects of LAP^+^ cell depletion on IFN-γ production.** Irradiated KDH-V cells (X-KDH-V) were inoculated into rats 21 days before inoculation of KDH-V cells. (A, B) DHP treatment was performed 12 or 14 days after inoculation of KDH-V cells. Blood cells were collected on day 17. (C, D) The DHP treatment was performed 11 days or 12 days after inoculation of KDH-V cells. Blood was collected on day 12. (E, F) DHP treatment was performed 11 days after inoculation of KDH-V cells. Blood was collected on day 14. (G, H) DHP treatment was performed 16 days after inoculation of KDH-V cells. Blood was collected on day 21. (B, D, F, H) The number of IFN-γ-producing cells in peripheral blood cells was determined by an ELISpot assay. Means and standard deviations of 6 to 8 wells of culture are shown.

**S2 Fig. Platelet count after DHP.** Column treatment was performed twice in three cancer rats. Squares and circles show platelet counts before and after DHP, respectively. The platelet counts decreased immediately after DHP (arrows). The platelet counts 5 to 7 days after the first DHP were higher than those before the first DHP.

## Acknowledgements

The authors would like to offer their sincere condolences at the passing of Dr. Yoshihiro Endo and express their gratitude to him for his substantial contributions to the present study. The authors would also like to thank Ms. Setsuko Fujita, Ms. Mihoko Kobayashi, and Ms. Naoko Kitagawa for their technical support. This work was partly supported by grants from the Japan Science and Technology Agency (JPMJSF23AT), the Center for Clinical and Translational Research of Kyushu University Hospital (A224), and the Japan Agency for Medical Research and Development (JP19im0502006h).

